# When random variation results in functional significance

**DOI:** 10.1101/2023.10.10.555393

**Authors:** Jacob Barfield, Patrick A. Kells, Shree Hari Gautam, Jingwen Li, Woodrow L. Shew

## Abstract

Many functional properties vary dramatically across neurons in cerebral cortex. Two fundamental goals of systems neuroscience are to determine which neurons execute which functions and how the different functional properties of a neuron are related. Often, it is assumed that if two properties are uncorrelated, then there is no important relationship to report. Here we show that this assumption can lead to wrong conclusions; functional segregation can emerge, by chance, due to random variation when that variation is distributed according to skewed, heavy-tailed distributions. We reinterpret the results we previously reported in Kells et al 2019 [1], showing that they are a prime example of functional segregation due to random variation.

## Introduction

As technological innovations push towards recordings of larger and larger populations of neurons (e.g. [2, 3]), it becomes feasible to assess similarities and differences across neurons with greater statistical rigor. Such assessments have revealed that many properties of neurons are not normally distributed across neurons [4]. Firing rates [5–7], synaptic strengths [5, 8, 9], pairwise spike correlations [10], dendritic spine size [11], and axon calibers [12] are a few examples of properties that seem to be distributed according asymmetric, skewed distributions with ‘heavy tails’. This means extreme outliers, although not common, occur far more often than one would expect for normally distributed quantities.

One tradition in data analysis is to ignore outliers, assuming that there is something wrong with the data points that fall far from the mean [13, 14]. However, with more samples collected, sometimes it turns out that there is nothing wrong with the outliers, rather the data is simply distributed in a skewed non-Gaussian, heavy-tailed way. In this case, ignoring the outliers can result in profoundly wrong interpretations. For example, considering synaptic efficacy; suppose we find that 1 in 1000 synapses is so strong that a single spike arriving at the presynaptic side can elicit an action potential from the post-synaptic neuron [5, 8, 9]. Considering that each neuron has 1000s of afferent and efferent synapses, this implies a ‘backbone’ network of strong synapses that can propogate signals very effectively without even considering the multitude of weaker, typical synapses [15] similar to the ‘rich club’ structure that has been observed at cellular scales [16] and at macro scales [17]. Thus, in the case of synaptic efficacy, ignoring the outliers may miss the most important and reliable circuit for transmitting information.

In this paper, we address another interesting implication and potential mistake when dealing with long-tailed distributions of neuronal properties. The case we consider is motivated by our previous study of neurons in motor cortex [1]. In this study a weak anti correlation was discovered between different forms of body coupling and population coupling. Some neurons seemed to be strongly correlated with body movement (i.e. strong body coupling), while a different population seemed to be strongly correlated with ongoing brain activity (i.e. strong body coupling). The anti-correlation was evidence that these two properties were functionally segregated, suggesting that neurons that are highly coupled to the movement of the body are not highly coupled to the activity of the population of neurons being studied and vice versa. However, the weakness of the anti-correlation cast some doubt on this possible type of functional segregation. In this paper we propose a new way to look at the data that allows this functional segregation to be analyzed irrespective of whether there is a strong, or even statistically significant, correlation between different properties.

## Materials and methods

### Animals

Experimental data from Kells et al [1] was used in this study. All procedures were carried out in accordance with the recommendations in the Guide for the Care and Use of Laboratory Animals of the National Institutes of Health and approved by University of Arkansas Institutional Animal Care and Use Committee (protocol #14048). We studied adult male rats (n = 6 Rattus Norvegicus, Sprague–Dawley outbred, Harlan Laboratories, TX, USA).

### Data analysis

#### Experimental Data Analysis

Three different forms of experimental data analysis were used in this study. Population coupling was defined as

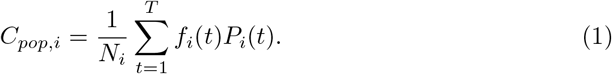

where *f*_*i*_(*t*) is the spike count in the *t*^*th*^ time bin of neuron *i, N*_*i*_ is the total number of spikes for neuron *i* over the whole recording, and the population spike count (for all but the *i*th neuron) is

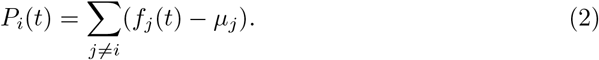

Where the mean spike count for neuron *j* is *µ*_*j*_. The population coupling is poorly estimated when there are only a small number of recorded neurons. For this reason, here we only considered recordings with more than 5 units.

Following Kells et al (2019), body coupling was measured in 2 different ways. Both methods were based of of the rat’s body movement and how strongly each recorded unit is related to this movement. The first way we assessed body coupling was based on a movement-triggered-average spike rate (MTASR) waveform. This is the average of the spike rate waveform for each neuron within a ±1*s* window around the onset or cessation of body movement. The value of body coupling *BC*_*M*_ was defined as the standard deviation of this MTASR waveform. The second property used to analyze body coupling is a spike-triggered-average body speed (STABS) waveform. This is the average of the body speed waveform within a ±1*s* window of each time a neuron fires. The body coupling *BC*_*S*_ is defined as the standard deviation of this STABS waveform for each neuron. In both cases the waveforms were low-pass filtered (1.5 Hz dutoff frequency) and then normalized by their mean.

#### Functional Segregation Procedure

Here we propose a test for functional segregation. For a given population of neurons with two functional properties measured for each neuron, let us call them A and B, we ask the question: Is the population functionally segregated into two sub-populations, one that does A and another that does B? To answer this question we suggest a new way to quantify functional segregation as shown in Eq (3).

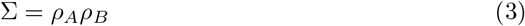

where *ρ*_*A*_ is the fraction of the range of A values greater than a threshold Θ_*A*_ and *ρ*_*B*_ is the fraction of the range of B values greater than a threshold Θ_*B*_. All possible pairs of thresholds are tested and then chosen to maximize Σ while meeting the constraint that no single neuron has *A >* Θ_*A*_ and *B >* Θ_*B*_ (the Θ thresholds are the right and top boundaries, respectively, of the red and blue shaded areas in Figs 1 and 2; the constraint enforces that there are no neurons in the white area in the upper right section of the plots). In addition to a measure of functional segregation we also developed a measure for how statistically significant these results are. Here we propose a measure for statistical significance that is the fraction of time a standard normal distribution with the same mean and standard deviation as the data is able to produce a larger value of functional segregation.

**Fig 1.**
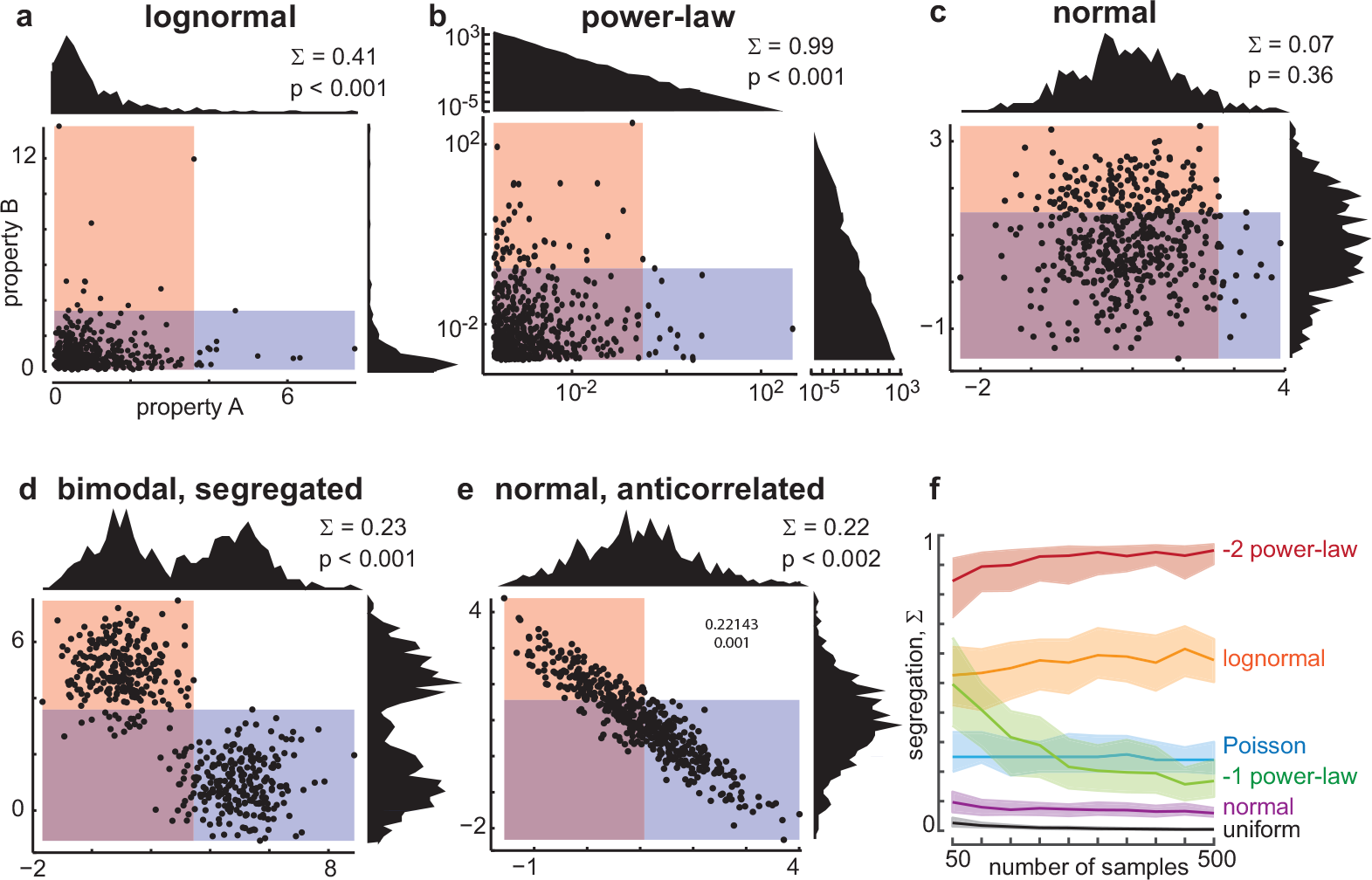
Functional segregation from random distributions. Σ quantifies functional segregation by chance and in more traditional scenarios. (a-e) Each point in the scatter plot represents one neuron. The horizontal and vertical coordinates of each point are determined by values for two hypothetical properties of the neuron (properties A and B, respectively). Distributions of property A are shown above each scatter plot. Distributions of property B are shown to the right of each scatter plot. The red shaded area indicates the range of A that is less than Θ_*A*_. The blue shaded area delineates B less than Θ_*B*_. The neurons in the red area that does not overlap the blue area are selective for property B and functionally segregated from the neurons in the non-overlapping blue area, which are selective for property A. Our segregation measure Σ quantifies the range of non-overlapping red and blue areas. Skewed distributions like the log-normal case (a) and the power-law (b) can exhibit significant Σ even though properties A and B are uncorrelated. Note that the power-law distributions and scatter plot have logarithmic axes for ease of visualization. Uncorrelated normally distributed properties are not significantly segregated (c). Anti-correlated properties also have significant segregation (d, e). f) The Σ measure is insensitive to sample size for more than a few hundred samples.

**Fig 2.**
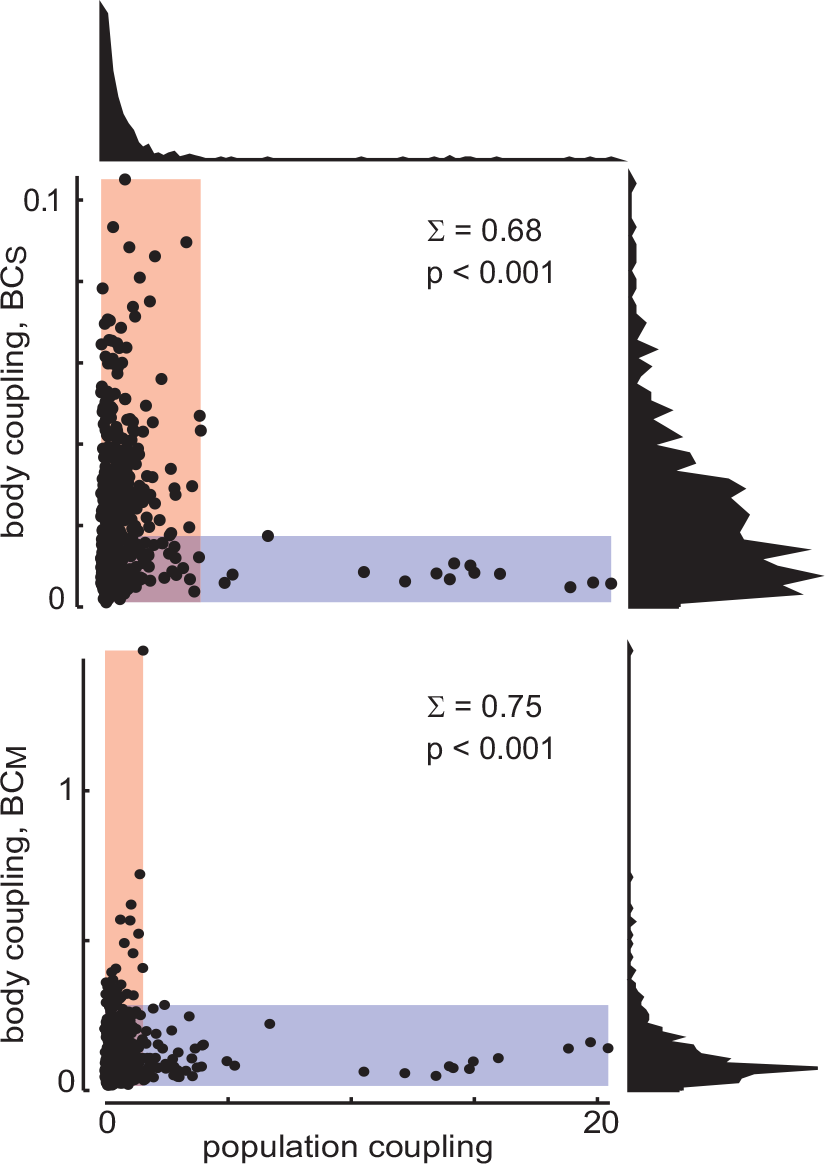
Functional segregation of population coupling and body coupling in motor cortex. Due to the skewed distributions of population coupling and body coupling, these two properties are strongly functionally segregated, even though these two properties are weakly correlated.

### Statistical Models

In this study we looked at numerous different statistical distributions of random numbers. These distributions range from the standard normal distribution to a power law distribution with exponent of -2 to an anti-correlated bi-modal distribution. Most of the distributions were made using a uniform distribution of random numbers and inverse transformation sampling was used to convert the uniform distribution into a custom distribution. Every distribution was run using 100 different seeds for the random number generators and were run with varying numbers of samples (500 samples each for the distributions in Fig 1a-e). For power law distributions and the uniform distribution we implemented upper and lower limits to ensure that the inverse transformation process was possible. The lower limit was chosen to be 0.01 and the upper limit was chosen to be 100. These same distributions did not have a way to set the mean and standard deviation so we implemented a tolerance level of 0.1 meaning that the distribution that was generated must have a mean and standard deviation that is withing 0.1 of the mean and standard deviation chosen for all of the rest of the distributions.

## Results & Discussion

In this study we consider two different functional properties that are governed by random distributions. We considered numerous different distributions for properties A and B and performed the above mentioned analysis on the data. These results are shown in Fig 1a-e. We also investigated how the number of samples effects the results of the functional segregation analysis (Fig 1f). We found that as the number of samples used increases the accuracy with which a value for Σ can be determined increases making the results more reliable. The results also show that for heavy tail distribution there is almost always a large value for functional segregation Σ which tends to increase slightly as the number of samples increases (with exception of the power law with exponent equal to -1). The distributions used that are not heavy tail all resulted in low values for functional segregation that remained relatively constant as the number of samples increased. We also found that beyond about 200 samples there was no significant change in the value for functional segregation as sample size increased.

There were two distributions analyzed that were not uncorrelated. These were the bi-modal segregated distribution and the normal anti-correlated distribution (Fig. 1d and Fig. 1e). Both distributions were made using a functional relationship that produces distributions that are anti-correlated. These cases were designed in such a way that they would produce a certain value of Σ but also be anti-correlated. The bi-modal distribution shows less correlation than the normal anti-correlated distribution.

In our previous study of rat motor cortex [1], which motivated our considerations here, we found that neurons in deep layers of rat primary motor cortex were functionally segregated, very similar to the scenario described here. The two properties we considered were population coupling and body coupling. The population coupling of a neuron quantifies how its firing covaries with the firing rate of the population in which it is embedded [1, 18]. The body coupling of a neuron quantifies how much its firing covaries with movements of the body [1]. We found that neurons with high body coupling had low population coupling and neurons with high population coupling had low body coupling (Fig 2a). This observation suggests that the neurons are functionally segregated into an ‘internal’ group (those with strong population coupling) and an ‘external’ group (those with strong body coupling). We did not observe overlap in these groups; there were no neurons with both very high body coupling and very high population coupling. This data was analyzed using this new framework for functional segregation, and the results of this analysis are shown in Fig 2. In these figures we show that for both forms of body coupling there is a significant value for Σ that arises when compared with population coupling. This tells us that the neurons that are strongly coupled to the spiking activity of the network are not strongly coupled to the motion of the body and vice versa. This is significant because it confirms one of the main conclusions from the Kells et al. paper without relying on the use of correlation analysis. This result also shows that there is significant functional segregation present in these systems and that this result is very statistically significant with a p value *<* 0.001. In the original paper this result relied on a weak peaked but statistically significant relationship between the two properties that showed a weak correlation between the two, but the properties were nearly uncorrelated. We believe that this functional significance being significant while the correlation between the properties is almost non existent reflects a real and relevant finding that functional segregation should not rely on the existence of a correlation.

## Conclusion

In this paper we proposed a new method for analyzing functional segregation that accounts for properties that are not Gaussian distributed and does not rely on traditional methods of measuring correlation between properties. We have shown that functional segregation can arise by chance from random variations across neurons. This is most evident when the segregated properties are are drawn from heavy tail distributions, which is important because heavy tail distributions are common when considering the properties of real neurons. In addition our findings expand upon similar ideas in which a randomly wired network of neurons can do quite useful things. This idea manifests in liquid state compouting [19] and the ‘preconfigured brain’ concept proposed by Buzsaki and colleagues [4, 5].

## Acknowledgments

This work was funded in part by the Foundational Questions Institute and Arkansas Biosciences Institute. The experimental data shown was first published in Kells et al. Nature Communications 2019 [1].

